# Unique Cancer Motility Behaviors in Confined Spaces of Microgroove Topography with Acute Wall Angles

**DOI:** 10.1101/694638

**Authors:** T. Yaginuma, K. Kushiro, M. Takai

## Abstract

In recent years, many types of micro-engineered platform have been fabricated to investigate the influences of surrounding microenvironments on cell migration. Previous researches demonstrate that microgroove-based topographies can influence cell motilities of normal and cancerous cells differently. In this paper, the microgroove wall angle is altered from obtuse to acute angles and the resulting differences in the responses of normal and cancer cells are investigated to explore the geometrical characteristics that can efficiently distinguish normal and cancer cells. Interestingly, trends in cell motilities of normal and cancer cells as the wall angles are varied between 60-120° were different, and in particular, invasive cancer cells exhibit a unique, oscillatory migratory behavior. Results from the immunostaining of cell mechanotransduction components suggest that this difference stems from directional extension and adhesion behaviors of each cell type. In addition, the specific behaviors of invasive cancer cells are found to be dependent on the myosin II activity, and modulating the activity can revert cancerous behaviors to normal ones. These novel findings on the interactions of acute angle walls and cancer cell migration provide a new perspective on cancer metastasis and additional strategies via microstructure geometries for the manipulations of cell behaviors in microscale biodevices.

**Statement of Significance:** Cancer metastasis is the leading cause of cancer patient deaths, and yet how the microstructures within the body affect this cell migration phenomenon is not well understood. In this paper, microdevices containing microgroove structures of varying geometries, in particular obtuse and acute angles, were utilized to monitor cell motilities of various cancer cells to understand the influences of the geometrical features of microstructures on cancer metastasis. Surprisingly, it was found that the acute angle geometries lowered the persistence of migration for cancer cells, which was a totally different response from non-cancerous cells. These new findings would enable the next-generation biodevices to analyze, separate and capture cancer cells, as well as shed light onto the underlying mechanisms of cancer metastasis.

## 1. Introduction

Cells in the body are constantly interacting with the surrounding microenvironments such as the extracellular matrix (ECM) and other cells. Depending on the conditions of such microenvironments, cells are known to alter their functions including adhesion (1-3), migration (4-6), differentiation (7), etc. Specifically, cell migration is one of the most important cell functions that plays an important role in various physiological phenomena, such as immune response (8), tissue formation (9-11), and cancer metastasis (12-14). The interactions between migrating cells and the surrounding environment are extremely complicated, so in order to simplify and isolate such interactions, many types of analytical platforms have been fabricated and the influences of surrounding microenvironments on cell migration have been investigated by utilizing these platforms. These studies have reported that cell migration is affected by both chemical and physical environments such as surrounding chemical gradients, surface chemistry, stiffness and surface topography (4, 5, 15-22).

Conventionally, the above studies have been conducted on 2D substrates. However, in recent years it has been found that the microscale three-dimensional topography on the substrate surfaces could induce a unique behavior of cells different from the 2D culture conditions, and furthermore drastically alter the cell motility (1, 14, 23-29). Moreover, it has been found that the degree of influence of 3D topographies is different depending on the capability of each cell to sense and interact with the substrate material. The invasiveness of breast cancer cells was markedly enhanced in 3D culture methods compared to conventional 2D culture methods, while tumorigenic cancer cells and normal cells did not show the invasion in the same matrix (30). The microfibrillar pattern mimicking the extracellular matrix morphology induced the chemotaxis of specific brain cancer cells, which was not observed on the 2D substrates (31). Across these studies, invasive cancer cells were found to behave distinctively by being trapped in a 3D microtopography. Depending on the surface properties of the surrounding 3D microtopographies, such as cell adhesiveness, pore size, and stiffness, they exhibited different migratory modes (14, 27). Lamellipodium migration, lobopodium migraion and amoeboid migration are representative migratory modes based on different migration principles. In other words, the confinement of the certain 3D microtopographies was found to induce such modes of cell migration, in a different manner from the macroscopic 3D matrices or 2D substrates. In addition, as the previous researches have demonstrated, cells can dramatically change their migratory behaviors according to the surrounding microscale topography, based on the property of each cell type. These researches on the regulation of cell migration utilizing 3D topographies are crucial in not only understanding both the fundamental properties of cells and various phenomena in the body, but also to provide the foundation for new technologies for the manipulation or separation of cells, which will lead to further applications for the diagnosis or therapy (18, 31, 32).

As one potential uses of substrate topographies in regulating cell migration, we have previously reported that microgroove-based topographies at the size scale of cells enhanced the cell motility and emphasize the motility differences between normal and cancer cells (33, 34). In addition, for normal cells, the topographical effects on cell migration were found to be more remarkable along 90° walls than >90° ones. It was suggested that this simple system combining 2D bottom planes and wall planes partially induced certain cell migration modes on 3D microtopographies, and the wall angle was a key factor. Therefore, in this research, the microgroove wall angle was altered from the range of obtuse to acute angles, and the differences in the responses of several types of cells including metastatic cancer cells to the microgroove topographies with different wall angles were investigated. Especially, the acute angle walls could physically confine cells like 3D-culture and were expected to induce a unique cell migration modes in between 3D microtopographies and 2D flat surfaces. These studies would allow us to deepen our understandings of the effects of microgroove topographies on cell migration and lead to more effective cell regulation. Furthermore, it may lead to new insights into the fundamental cellular mechanisms in a physically confined space and may open up different avenues in utilizing microscale platforms for cell behavior analysis.

## 2. Results & Discussions

### 2.1. Fabrication of PDMS microgroove structure with various wall angles

PDMS substrates with microgroove topographies with different wall angles were fabricated by replica molding (**Figure 1**). To assess the fabricated topographies, bovine serum albumin (BSA) tagged with fluorescence was adsorbed on the surface of the substrates and the fabricated topographies were observed using a confocal microscope. From the 3D confocal imaging of the fluorescently labeled BSA, both vertical and slanted microgroove structures were confirmed to be successfully fabricated. As the images of cross-sections show, the slanted microgroove had acute edges on one side and obtuse edges on the other side. By utilizing microgroove Ni-molds with different wall angles by 10 degrees in three steps from right angle, microgrooves with 7 different wall angles (60, 70, 80, 90, 100, 110, and 120 degrees) were prepared for cell experiments.

**Figure 1.**
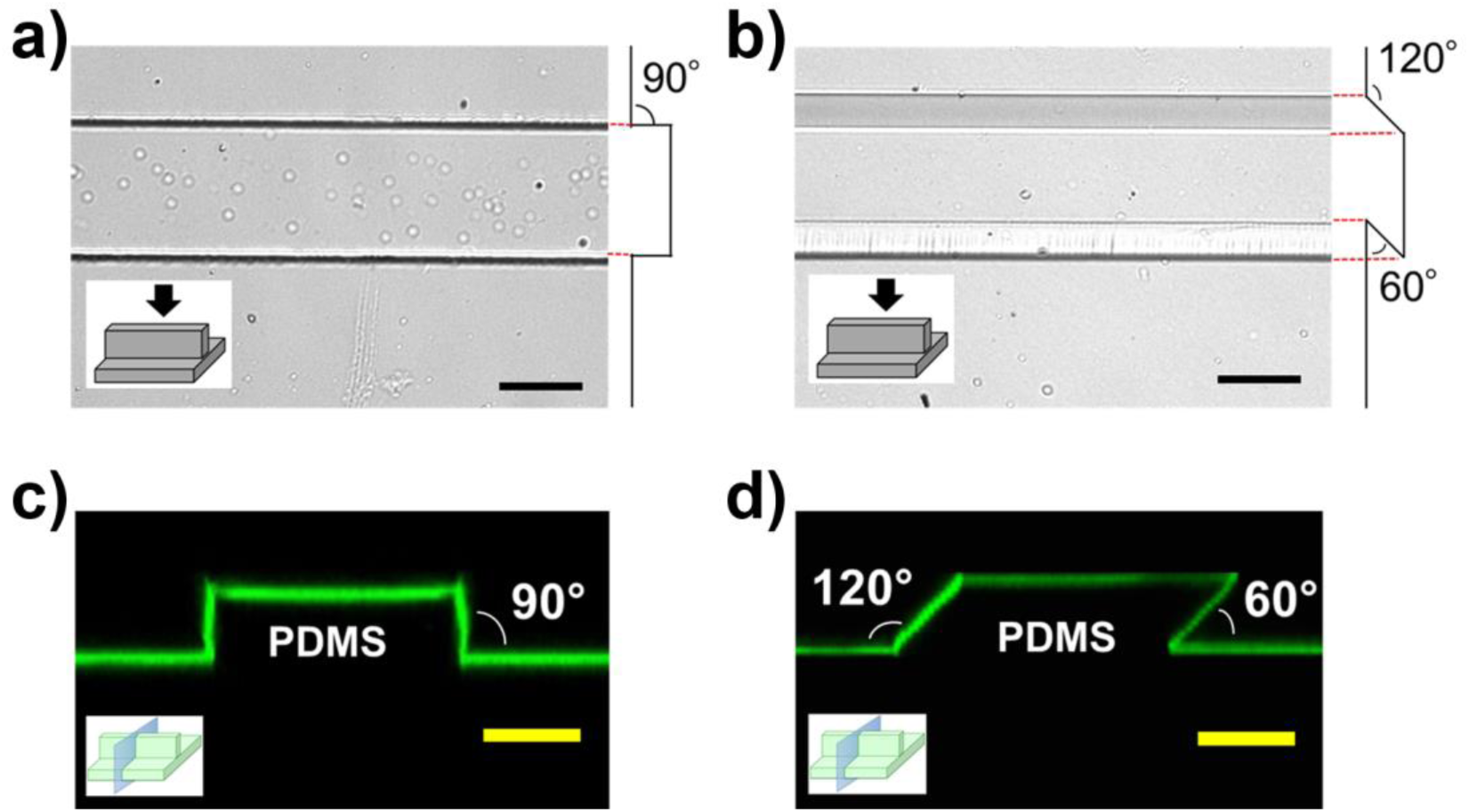
Bright-field image of the microgroove-based topographies with (a) a vertical wall and (b) a slanted wall. Confocal images of the cross sections of the microgroove-based topographies with (c) vertical wall and (d) slanted wall visualized by the adsorbed FITC-BSA. (Scale bar, 50 μm)

### 2.2. Cell motility on microgroove structures with various wall angles

Normal MCF10A cells, non-invasive cancer MCF7 cells and invasive cancer MDAMB231 cells were seeded on the substrates with microgroove structures of different wall angles from 60 to 120°. Cell motilities on the microgroove topographies with different angles were evaluated from time-lapse imaging (**Figure 2**). These were evaluated as 2 indices: velocity and persistence. The persistence was an index for the directionality and was evaluated by measuring the distance that cells migrated until they changed their direction by >90 degrees. According to this definition, for cells traveling along the microgroove structure, the absolute value of the distance traveled unidirectionally along the microgroove was measured as the persistence length.

**Figure 2.**
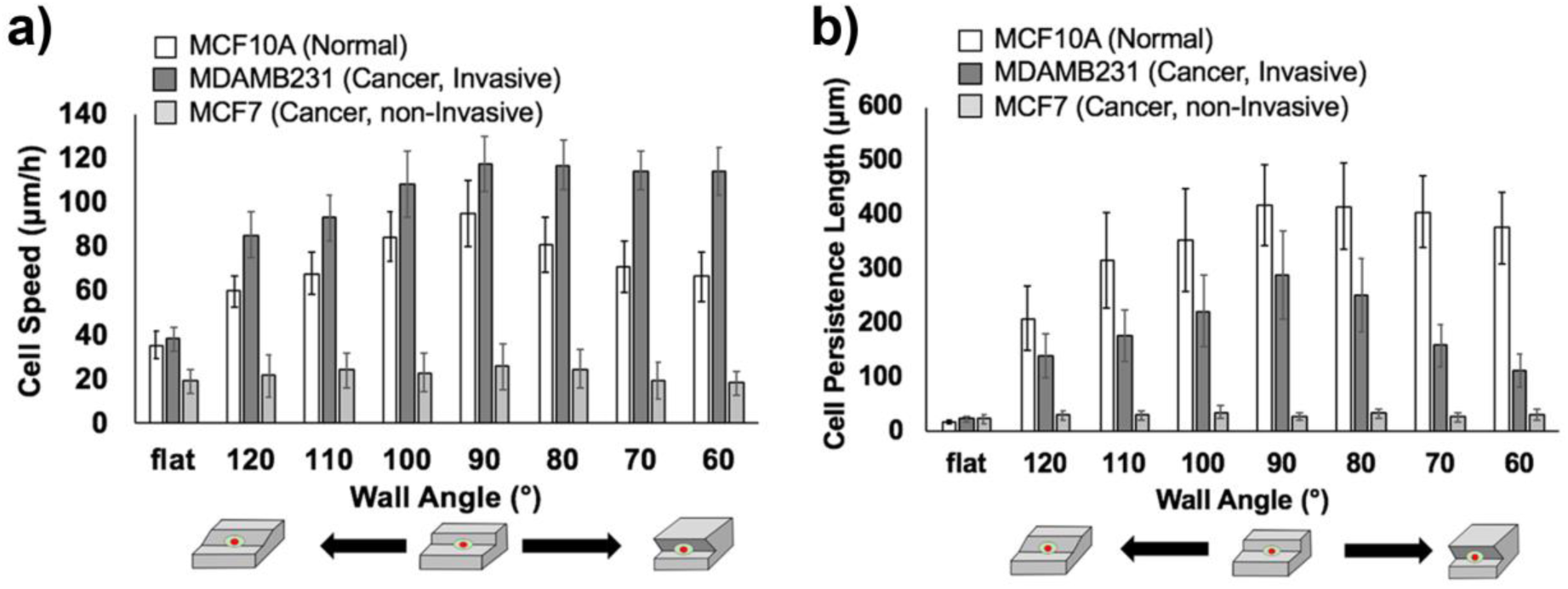
(a) Cell velocity and (b) cell persistence length of MCF10A cells (normal), MDAMB231 cells (cancer, invasive) and MCF7 cells (cancer, non-invasive) along the edges of microgroove topographies with 60 to 120° walls and on the flat surface.

From the results in Figure 2, it was apparent that each cell type exhibited different responses to the microgroove topographies with varying wall angles. First, for normal cells and invasive cancer cells, there was no significant difference between the motilities of these cells on a flat surface (normal cells: velocity 36 ± 6 μm/h, persistence18 ± 2 μm; invasive cancer cells: velocity 38 ± 6 μm/h, persistence 23 ± 4 μm). For 120-90° angles, normal and cancer cell motilities showed similar trends, where the velocities and persistence lengths of both types of cells increased as the wall angle decreased (normal cells: from 60 ± 7 to 95 ± 15 μm/h for velocity, from 210 ± 60 to 419 ± 74 μm for persistence length; cancer cells: from 85 ± 10 to 118 ± 12 μm/h for velocity, from 141 ± 41 to 290 ± 81 μm for persistence length) (Supporting Material; **Movies S1-S4**). These results are consistent with our previous findings that cells along obtuse walls had difficulties in arranging the cytoskeleton along the microgroove edges, which led to the decrease of the motility parameter (33, 34). But when the wall angle was 60-90°, different trends were observed between normal and cancer cell motilities (Supporting Material; **Movies S5**,**6**). In terms of cell velocity, the velocity of normal cells became smaller as the wall angle changed from 90 to 60° (from 95 ± 15 to 66 ± 11 μm/h), while the velocity of cancer cells did not change (from 118 ± 12 to 114 ± 11 μm/h), though it was overall larger than that of normal cells. In terms of persistence, the persistence length of normal cells did not change significantly as the wall angle changed from 90 to 60° (from 419 ± 74 to 377 ± 65 μm), but the persistence length of cancer cells decreased (from 290 ±81 to 113 ± 31 μm) and was smaller than that of normal cells overall. Regarding the persistence length of the cells, there was a characteristic difference in migration behaviors for each cell type. While normal cells continued unidirectional migration along the walls of microgroove topographies until they collided with other cells or entered the mitotic phase, cancer cells frequently reversed direction and thus oscillated along the wall, which was reflected on the decrease in persistence lengths (Supporting Material; Movies S5, S6).

For non-invasive cancer cells, motility did not change, regardless of the variations in the wall angle or the presence of the topography (velocity: 18 ± 5 μm/h to 25 ± 10 μm/h, persistence length: 22 ± 9 μm to 36 ± 12 μm). It seemed to be caused by the lack of the 3D geometric recognition abilities of non-invasive cancer cells, which might be derived from the decreased interactions with underlying substrates compared to the other cell types (Supporting Material; **Movie S7**) (34).

In addition to the motility parameters and related to the recognition of the 3D topography, there was a significant difference in the morphology of each cell type in response to various geometries. By evaluating the ratios of cells extending along the microgroove structure with unilamellar morphology (motile cells), it was found that there was a large difference between the normal and cancer cell populations (Supporting Material; **Figure S1**). Along the vertical angle wall, about 70% of normal cells adopted the motile morphology but for 2 types of cancer cells the percentage of motile cells greatly decreased compared to the normal cells. This result indicated that 3D geometric recognition abilities of normal cells are much higher than cancer cells. Moreover, these results, in which invasive cancer cells with motile morphologies showed drastically higher motilities than non-invasive cancer cells, could reflect one of the heterogeneous features of tumors in which some cells display significantly higher motility over others, even though the average motility of the entire cell population is low due to the existence of the many non-motile cells (13, 14, 34). At the same time, it was also demonstrated that the microgroove systems may be useful in differentiating the motile and non-motile cancer cells even in a heterogeneous cancer population.

Lastly, invasive cancer cells and normal cells showed similar trends in motility parameter when along microgroove topographies with obtuse and vertical angle walls (120-90°), but they showed completely different trends when along the microgroove topographies with acute angle walls (90-60°). In particular, invasive cancer cells showed the unique frequent reversal behaviors, and thus when along the acute angle walls, invasive cancer cells showed the migration behavior that can be clearly distinguished from other two cell types. These findings suggested that the acute wall angles could influence the cell migration more and induce the unique behaviors of different cell types.

### 2.3. Immunostaining of cells and the analyses of extension behaviors and focal adhesions

To investigate the differences of cell motilities along acute angle walls, especially that between the normal (MCF10A) and invasive cancer cells (MDAMD231), key cell mechanotransduction components were visualized by immunostaining. The actin stress fibers were chosen to visualize the cytoskeletal reorganization in response to the microstructures, while vinculin was chosen to visualize the focal adhesions (FAs) to understand the cell migration direction and persistence. Consistent with the motility evaluations, we found that the extension behaviors and adhesion states were signficantly different between normal and invasive cancer cells along the acute angle walls.

First, the morphologies of the cells along the 90-60° walls were evaluated with confocal microscopy after staining the nucleus and actin filaments (**Figure 3**). For both normal and invasive cancer cells, the extensions in the direction parallel to the wall were enhanced by the acute angle walls compared to the vertical walls. However, upon observing the cross sections of these cells, there was a striking difference in the manner of extension. For normal cells, as the wall angle decreased from 90° to 60°, those extensions in different directions of movement such as upward or lateral to the acute angle wall were enhanced due to the increased confinement (Figure 3a,b). Meanwhile cancer cells further extended only in the direction of migration along the 90-60° walls (Figure 3c,d). This difference in the extension behavior could explain the cause of the decrease in cell velocity of normal cell along the <90° walls and corroborate the fact that the velocity of invasive cancer cells was independent of the wall angle. As for the non-invasive cancer cells (MCF7), even along the wall of the microgroove topography, actin filaments were not aligned along the wall, unlike the other two cell types (Supporting Material; **Figure S2**). This result is consistent with the result that the motilities of non-invasive cancer cells did not change by the presence of the microtopography.

**Figure 3.**
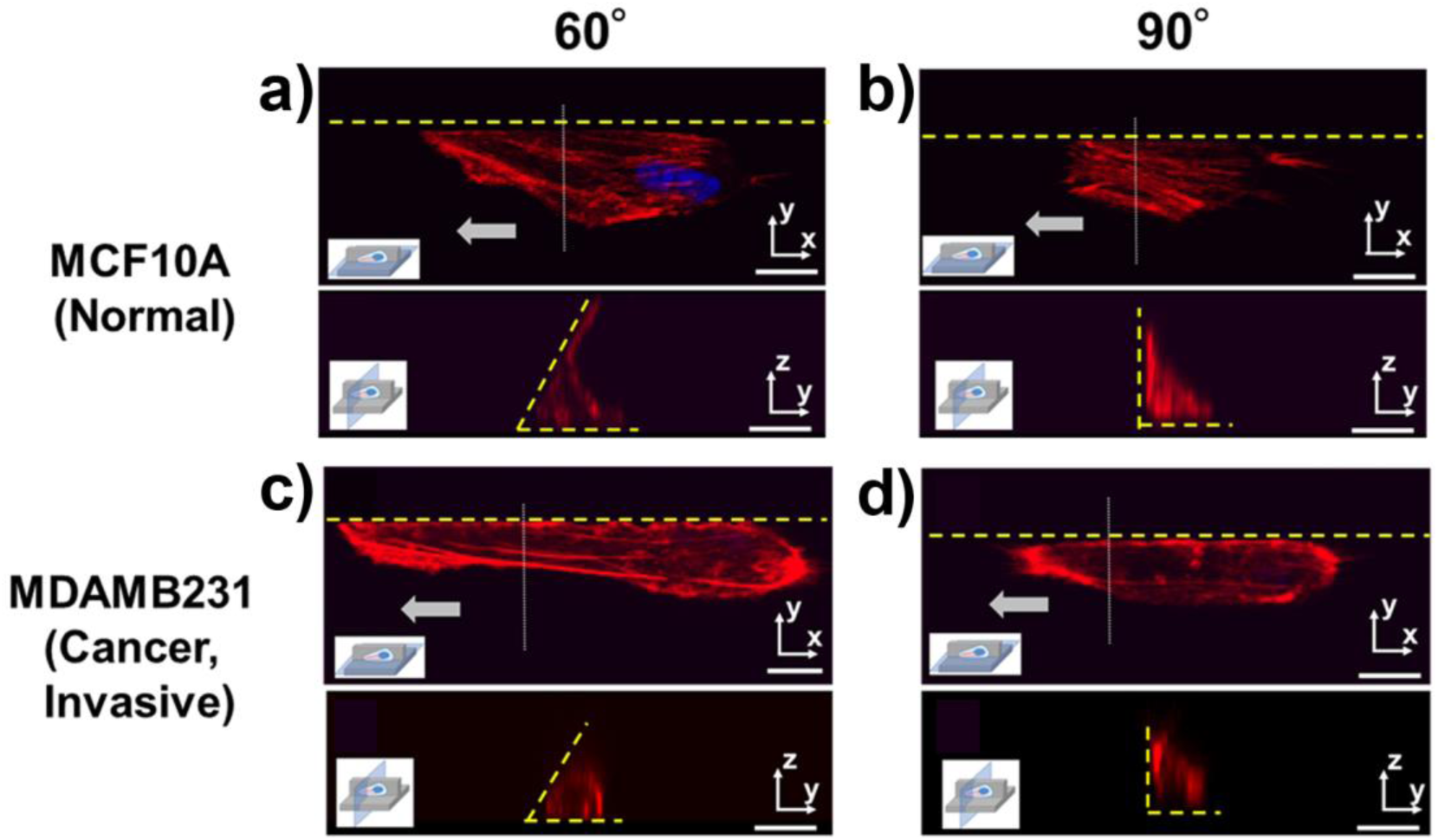
Immunostaining (63x) of actin (red) and nucleus (blue) of (a,b) MCF10A (normal) and (c,d) MDAMB231 (cancer, invasive) cells along the edge of microgroove topographies with 90° and 60° walls, with the same cell visualized on the X-Y planes and the Y-Z planes. Yellow dotted lines represent the microgroove wall boundaries. Gray arrows represent the direction of cell migration. (Scale bar: 10 μm)

Then to evaluate the adhesion states of cells, vinculin molecules were stained and also quantified through confocal microscopy images (**Figure 4**). Vinculin is one of the main components of FAs, which represent the strength and directionality of cell adhesion.^[19, 35]^ For normal cells, FAs along the 60° and 90° walls and on the flat surface were found to be densely concentrated at the leading edge of the migrating cell (Figure 4a,b). Those of invasive cancer cells along the flat surface showed a similar trend to the normal cells. However, regarding invasive cancer cells along the 60° and 90° walls, FAs appeared densely at the trailing edges as well as at the leading edges and that trend was more enhanced when the wall angle became smaller (Figure 4c,d).

**Figure 4.**
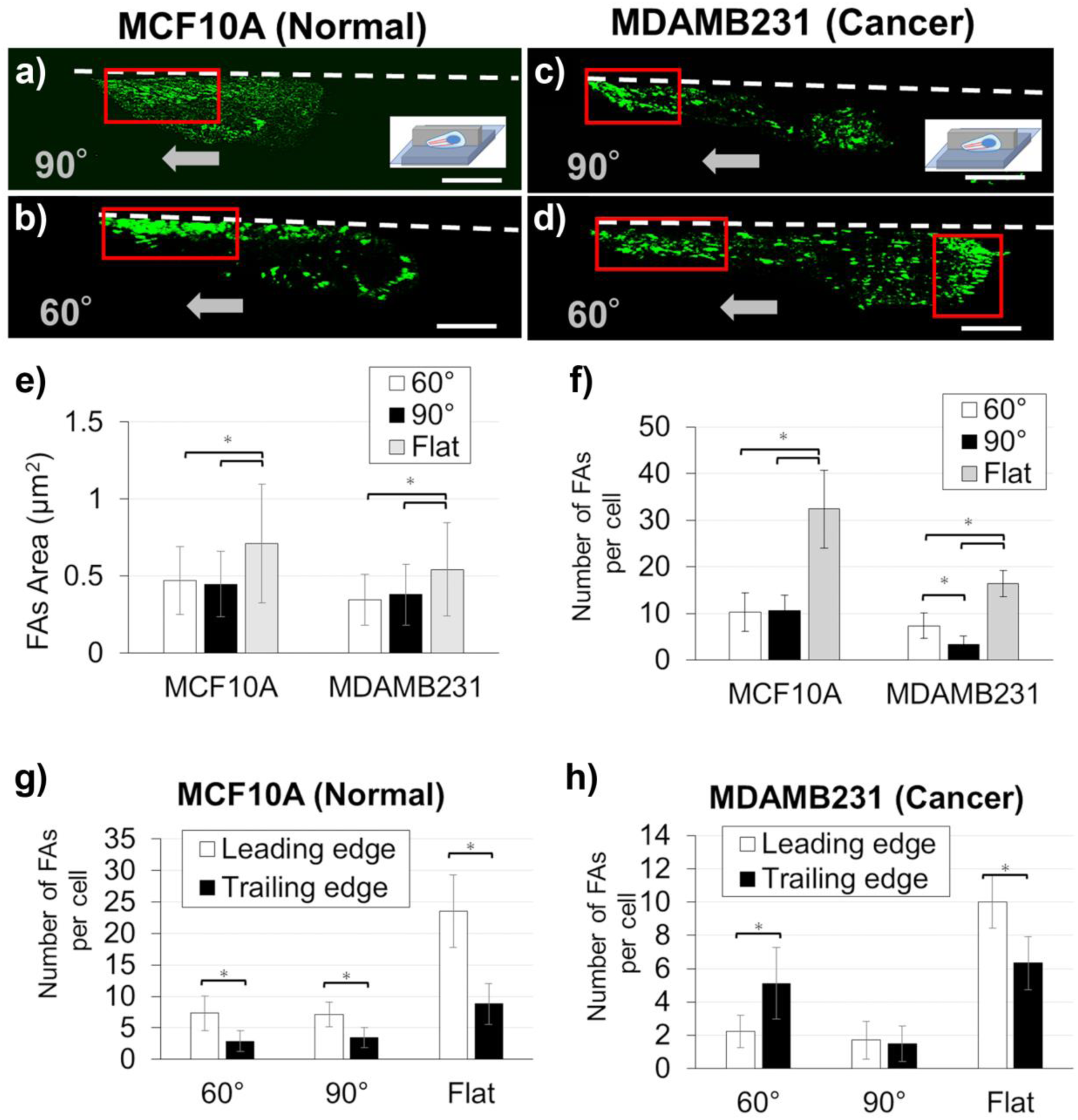
Immunostaining (63x) of vinculin (green) of (a,b)MCF10A (normal cells) and (c,d) MDAMB231 (cancer cells) along the edge of microgroove topographies with 90° and 60° walls. White dotted lines represent the microgroove wall boundaries. Gray arrows represent the direction of cell migration. Red boxes indicate dense areas of focal adhesions. Quantitative image analyses of focal adhesions for MCF10A (normal) and MDAMB231 (cancer, invasive) along the edge of microgroove topographies with 90° and 60° walls, and on the flat surface were performed. (e) Focal adhesion area and (f) the number of the matured focal adhesions (>0.5 μm^2^) contained in each single cell are shown. The number of the matured focal adhesions (>0.5 μm^2^) contained in the leading edges and the trailing edges of (g) normal and (h) cancerous cells are shown. (Scale bar: 10 μm) (* : p < 0.05)

The FA properties, such as the number or the area of FAs, were also analyzed quantitatively from the obtained confocal images using ImageJ. In addition to the number of FAs, the distribution of FAs between the leading edges and the trailing edges of the cells were analyzed, in which each cell was divided in half and FAs were counted for each side. The results indicated that, for the area of FAs, there was no significant difference between the cells along the 60° and 90° walls, although FAs of normal cells were overall larger than ones of invasive cancer cells (Figure 4e). Regarding the total number of FAs, the trends in the FA number (Figure 4f) seemed to inversely correlate with the trends in persistence length (Figure 2b), suggesting that FA number is indicative of the persistence in cell migration. Taken together, these results also suggested that the wall angle influence the number of FAs, rather than the area of FAs. As for the distribution of FAs, for normal cells, the number of FAs at the leading edges were more than twice as many as that at the trailing edges (Figure 4g), consistent with their high persistence lengths. In terms of invasive cancer cells, although the number of FAs at the leading edges were more than that at the trailing edges on the flat surface, FAs formed equally at leading and trailing edges along the 90° walls (Figure 4h).

Moreover, along the 60° walls the number of FAs at the trailing edges were significantly larger than at the leading edges, which may reflect the underlying FA reversal preceding the reversal of cell migration direction. This unique underlying adhesion states of invasive cancer cells in the confined space of the acute angle wall explain the frequent reversal of the migration directions, which cause the drastic decrease of the persistence length of invasive cancer cells along the acute angle walls. At the same time, these findings suggest that acute angle walls may be a useful geometrical feature to trap only the invasive cancers in place.

### 2.4. Roles of myosin II in regulating cell motilities along acute angle walls

In order to prod the system further, the underlying mechanisms related to the difference in the motilities of normal and invasive cancer cells along acute angle walls were investigated. In particular, the focus was on the unique migratory mode of invasive cancer cells along acute angle walls from the perspective of migration machinery. The motilities of invasive cancer cells have been known to be greatly influenced by surrounding environments, and a physical confinement is one of the key factors influencing the cancer cell motility (14, 27). In a similar manner, microgroove topographies with acute wall angles may be perceived like a confined space by cells, possibly inducing the unique behavior of invasive cancer cells. Thus, factors relating to the cell migration in the confined space were investigated. In particular, myosin II has been known to be deeply involved in cancer cell migration within confined spaces and such cancer cell migration has been reported to be mainly driven by the contraction force generated by myosin II (14, 27, 36-38). In other words, the activity of myosin, which has been correlated to the degree of the physical confinement felt by cells, may be considered as a tunable parameter to regulate the mode of motility. Using blebbistatin, a myosin II inhibitor, the relationship between the cell migration mode along the acute angle walls and myosin II activity was investigated. Normal cell (MCF10A) and invasive cancer cell (MDAMB 231) were seeded on the substrate with the microgroove topographies and treated with blebbistatin (concentration in growth medium: 0 μм, 2.5 μм, 5 μм) (Supporting Material; **Figures 5, S3**).

**Figure 5.**
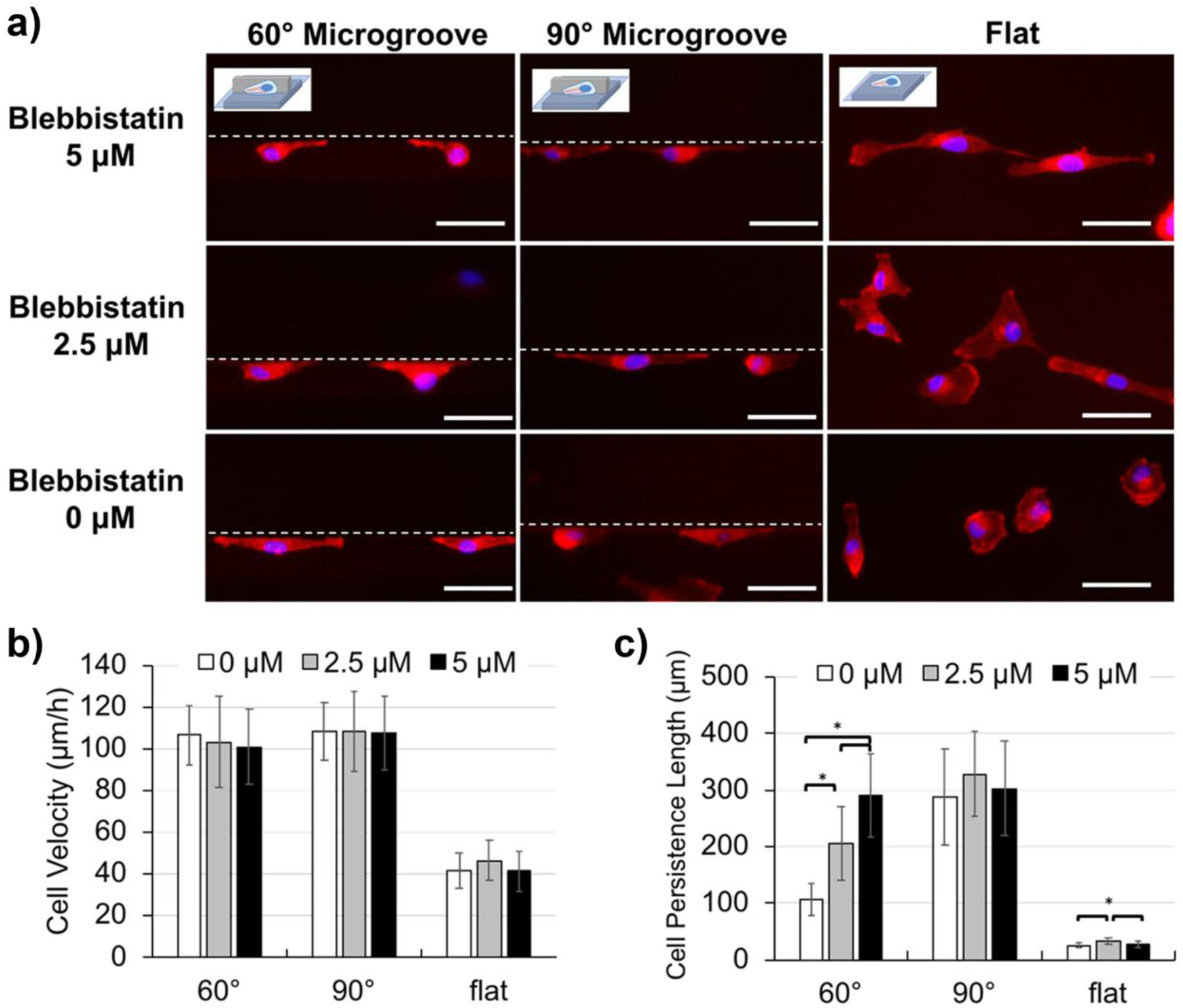
The morphologies and motilities of MDAMB231 cells (cancer, invasive) treated with blebbistatin, myosin II inhibitor. (a) Cell morphologies visualized by immunostaining (20x) of actin (red) and nucleus (blue) along the edge of microgroove topographies with 90° and 60° walls, and on the fat surface. Blebbistatin concentration was varied from 0 μм to 5 μм. White dotted lines represent the microgroove wall boundaries. (b) Cell velocity and (c) cell persistence length of MDAMB231 cells treated with blebbistatin. (Scale bar: 50 μm) (* : p < 0.05)

First the cell morphologies were observed and it was found that blebbistatin treatment greatly changed cell morphology (Supporting Material; Figure 5a, S3a). With respect to normal cells, the cells spread broadly on a flat surface with blebbistatin treatment. When these cells were along the microgroove topography, the lamellipodial extensions became larger and the detachment of FAs at the rear end of the cells seemed less efficient. There was no remarkable difference between the normal cells along 60° and 90° walls. Next, for the invasive cancer cells, the cells spread broadly on a flat surface with blebbistatin treatment, similar to the normal cells. When invasive cancer cells were along the vertical walls, although there was no significant change in the morphology, FA detachment also seemed less efficient at the rear end of the cells. In contrary, however, when along the acute angle wall, the bidirectional stretching behavior was suppressed by the blebbistatin treatment and the cells extended unidirectionally along the wall, which was the morphology associated with high motility (Figure 5a).

Quantitative evaluation of cell motilities under the influence of blebbistatin was also performed for normal and invasive cancer cells along 60° and 90° walls. First, for normal cells, blebbistatin treatment was found to greatly enhance both the cell velocity and the persistence length. This trend was observed commonly in all conditions regardless of the presence or wall angles of microgroove topography. Interestingly, the motility parameters were more enhanced when the concentration of blebbistatin was 2.5 μм than 5 μм (Figure S3b,c). To discuss the relationship between mobility of normal cells and blebbistatin treatment, the contribution of myosin II to normal cell migration should be considered. Generally, myosin II is well known to deeply contribute to cell contractility, but indirectly it is also involved in various mechanisms of cell migration, such as the formation of FA or the formation of actin stress fibers. Furthermore, suppression of myosin II has been reported to upregulate Rac1, which is a signaling factor that controls lamellipodial activity, and the activation of Rac1 is generally known to enhance cell motilities (39, 40). The motility enhancement by Rac1 activation is universal to the migration principle of normal cells and is consistent with the fact that the influence of blebbistatin treatment appeared similarly in all conditions regardless of the presence or the wall angle of microgroove topography. Furthermore, the morphology and the biphasic effects of blebbistatin on cell motility also suggest that at high concentrations, there may be not enough contraction to detach the FAs.

Next, the invasive cancer cells exhibited a different response to blebbistatin treatment compared to normal cells. It was found that the frequent reversal behaviors of invasive cancer cells along acute angle walls were neutralized by blebbistatin treatment (Supporting Material; **Movie S8**). This strongly suggested that the activity of myosin II may be deeply involved with the reversal behavior along acute walls. As a result of quantitative evaluation, it was found that the cell persistence length along the 60 ° wall increased as the blebbistatin concentration became higher (0 μм: 106 ± 28 μm; 2.5 μм: 206 ± 65 μm; 5 μм: 291 ± 73 μm), and interestingly, the influence of blebbistatin treatment did not appear in other geometrical conditions (Figure. 5b,c). This was a major difference between normal and invasive cancer cells, where for normal cells, the blebbistatin treatment enhanced the motility parameters at all conditions, while for invasive cancer cells, the influence of blebbistatin treatment appeared only in the persistence along the acute angle wall. This suggests that signaling and thus the mode of motility of invasive cancer cells along acute walls are indeed different from other conditions. In fact, several previous studies have reported that the aggregation and activation of myosin II were enhanced at the trailing edges of invasive cancer cells when they are trapped in spaces with strong physical confinement, such as non-degradable gels (14, 36-38). Taken together, the findings point to a similar mechanism of cell migration for acute angle walls, and it may be said that the physical confinement due to the acute angle wall promoted the activation of myosin II at the trailing edges of invasive cancer cells, which caused the frequent reversal behaviors. Moreover, the findings unraveled the modulation of myosin II as a potential strategy to undo cancer-specific migration behaviors in physical confinements.

## 3. Conclusion

In this paper, the relationship between the microgroove topography effects on cell migration and the wall angle was investigated. In summary, the differences among normal cells, non-invasive cancer cells, and invasive cancer cells became even greater when the wall angle of microgroove was in acute range than the wall angle was in obtuse range. We found that each cell type showed specific responses to the acute angle wall. Specifically, invasive cancer cells displayed a unique mode of motility where they frequently reversed the migration direction along the acute angle walls, while normal cells along the acute angle wall showed the similar migration behavior to the vertical wall but with only the velocity decreased. The non-invasive cancer cells did not get affected by the topographies, similar to our previous studies. Immunostaining of actin fibers revealed the reason why cancer cells maintained the fast velocity along the acute angle wall, while normal cell velocity decreased. Moreover, immunostaining of vinculins revealed that the frequent reversal behaviors stemmed from the unique adhesion formations. These results also suggested that the lamellipodial extensions and the adhesion is a key driving force in topography-directed migration. Lastly, it was found that such unique behaviors of invasive cancer cells along the acute angle walls were extensively involved with the myosin II activities, with many similarities to cancer cells exposed to physical confinements. The findings from this study highlight the implications of the 3D confinement to cancer cell migration and deepen our understandings of the effects of microgroove topographies on cell migration, providing new geometrical tools and strategies to be applied to analysis and manipulation of cell behaviors in microscale platforms.

## 4. Experimental Section

### 4.1. Fabrication of PDMS microgroove structure with various wall angles

Microgroove structures (100-μm-wide and 40-μm-high; wall angles = 60, 70, 80, 90, 100, 110, and 120 degrees) were fabricated by casting polydimethylsiloxane (PDMS; Silpot 184 W/C; Dow Corning) onto the custom-made Ni-based mold, which was created in collaboration with Optnics Precision, Co. Briefly, silicon wafers were spin-coated with a negative photoresist (40-μm-thick), exposed to ultraviolet light at different angles through the chrome photomask, developed using a developer, sputtered with a conductive layer (300-nm-thick), and finally cast with an electroforming metal containing Ni to electroform a Ni-based mold (300-μm-thick) (Supporting Material; **Figure S4**). The PDMS elastomer base and the curing agent were mixed at the 10:1 weight ratio. PDMS substrates with microgroove structures were rendered cell adhesive by adsorbing fibronectin (Sigma) for 1 hr at 37 °C (10 μg/mL), and then blocked with 0.1% bovine serum albumin (Sigma) for 2 hr at 37 °C. To evaluate the fabricated substrates, PDMS substrates with microgroove topographies were immersed in fluorescein-isothiocyanate-conjugated bovine albumin (Sigma) solution (10 μg/mL) for 2 hr at 37 °C. Then surface topographies were observed using a confocal microscope (LSM510; Carl Zeiss) at 20× magnification.

### 4.2. Cell culture

The invasive breast cancer MDAMB231 cells, non-invasive tumorigenic breast cancer MCF7 cells, and normal mammary epithelial MCF10A cells were used. MCF10A cells were cultured in growth medium composed of Dulbecco’s modified Eagle’s medium/Ham’s F-12 containing HEPES and L-glutamine (DMEM/F12, Invitrogen) supplemented with 5% horse serum (Invitrogen), 1% penicillin/streptomycin (Invitrogen), 10 μg/mL insulin (Sigma), 0.5 μg/mL hydrocortisone (Sigma), 20 ng/mL EGF (Peprotech) and 0.1 μg/mL cholera toxin (Sigma) and maintained under humidified conditions at 37 °C and 5% CO_2_. MDAMB-231 and MCF7 cells were cultured in growth medium composed of Dulbecco’s modified Eagle’s medium (DMEM, Invitrogen), supplemented with 10% fetal bovine serum (Invitrogen) and 1% penicillin/streptomycin (Invitrogen) and maintained under humidified conditions at 37 °C and 5% CO_2_. Cells were passaged regularly by dissociating confluent monolayers with 0.25% trypsin-EDTA (Invitrogen). Cells were passaged at 1:4 in growth medium. Blebbistatin (Sigma) dissolved in dimethylsulfoxide (DMSO) was used at 5 μM in growth media for the myosin II inhibition experiments.

### 4.3. Time-lapse microscopy and motility quantification

Cells were seeded at 5 × 10^3^ cells/mL in growth medium on the micropatterned substrates at 5 × 10^3^ cells/mL at 37 °C and 5% CO_2_. After seeding for 24 hr, the incubated cells were imaged at 10x magnification every 5 min for 24 hr using CCM-1.4II D/C Cell Culture Monitoring System (ASTEC). In this system, the cells were maintained at 37 °C and 5% CO_2_ in a heated chamber with temperature and CO_2_ controller during time-lapse imaging. CCM Software 2.4.5.2 (ASTEC) was used for image processing. To track cell motility on micropatterned surfaces, the position of the leading edge of migrating cells was tracked using CCM and ImageJ software. Cell motility was evaluated by 2 indices: velocity and persistence. To calculate the velocity, the distance cells migrated for 0.5 hr was measured. The persistence was evaluated by measuring the distance that cells migrated until they changed their direction by >90 degrees. According to this rule, in terms of cells along the microgroove structure, the absolute value of the distance traveled unidirectionally along the microgroove was measured and used as persistence length.

For the evaluation of MCF10A and MDAMB231 cells motilities along the microgroove structure, isolated cells extending along the microgroove structure with unilamellar morphology and migrating independently of other cells were targeted. In terms of MCF7 cells, which did not form the unilamellar morphology, isolated cells along the microgroove structure were targeted, regardless of the morphology. For evaluation of the motilities on flat substrates, isolated cells extended at a position 100 μm or more away from the microgroove were targeted. The unpaired, two-tailed Student’s t-test was used for statistical analysis. Differences were considered significant at p < 0.05. All the statistical analysis was performed using corresponding functions in Microsoft Excel.

### 4.4. Immunostaining of nucleus, actin and vinculin

Cells were fixed on 35 mm glass-bottomed dish (Iwaki) using 4% formaldehyde solution (Sigma) for 20 min at room temperature, permealized using 0.2% Triton X-100 (Amersham Biosciences) for 10 min at 4 °C, blocked with 0.1% bovine serum albumin (Sigma) overnight at 4 °C, and then incubated with dyes and antibodies. 4’,6-Diamidino-2-phenylindole, dihydrochloride (DAPI; Invitrogen) was incubated for 10 min to visualize the cell nucleus, Alexa Fluor 594 phalloidin (Invitrogen) was incubated for 1 hr to visualize the actin filaments, and primary anti-vinculin antibody (Abcam) was incubated overnight then tagged with secondary Alexa Fluor 488 goat anti-mouse IgG antibody (Abcam) for 2 hr to visualize the vinculin molecules, respectively. The fixed cells were visualized using a confocal microscope (LSM510; Carl Zeiss) at 63× magnification. Quantitative analysis of FAs was done by using ImageJ software.

## Supporting Material

Supporting Material can be found online at

## Author Contributions

T.Y. performed research; T.Y. and K.K. analyzed data; K.K. and M.T. designed research; T.Y., K.K. and M.T. wrote manuscript.

## Acknowledgements

This research was funded by the Grant-in-Aid for Young Scientists (A) (#16H05972) from the Japan Society for the Promotion of Science (JSPS). Lastly, we would like to thank Optnics Precision Co. for providing us with the mold with slanted features.

## Table of Contents (ToC)

### The table of contents entry

**Microgroove topography with acute angle wall** elicit different responses from normal and invasive cancer cells, in which normal cells migrate persistently, while cancer cells enter an oscillatory mode of migration. The findings shed light on the mechanisms underlying cancer migration in physical confinements and introduce microstructure geometries as tools to manipulate cell migration in microdevices.

#### ToC Figure

(55 mm broad × 50 mm high):

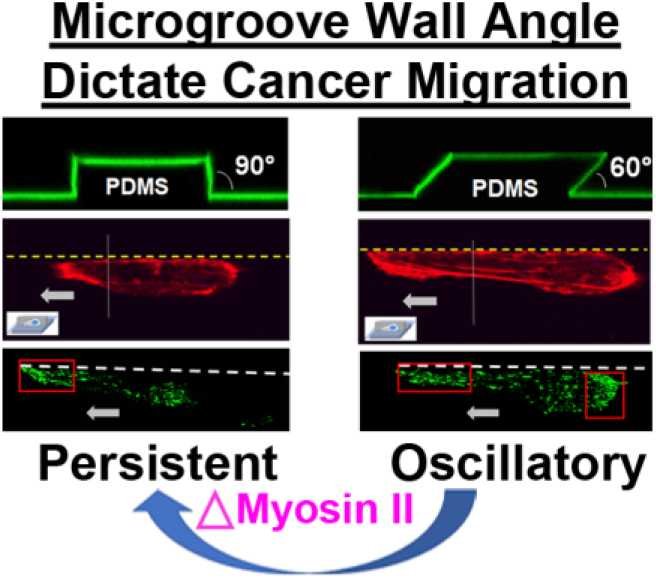

